# Mechanosensitive recruitment of Vinculin maintains junction integrity and barrier function at epithelial tricellular junctions

**DOI:** 10.1101/2023.09.08.556899

**Authors:** Lotte van den Goor, Jolene Iseler, Katherine Koning, Ann L. Miller

**Affiliations:** Department of Molecular, Cellular, and Developmental Biology; University of Michigan; Ann Arbor, Michigan, 48109; USA; Cellular and Molecular Biology Graduate Program, University of Michigan; Ann Arbor, Michigan, 48109; USA

**Keywords:** tricellular junctions, adherens junction, actin, Vinculin, tension, epithelium, barrier function, *Xenopus*, live microscopy, optogenetics

## Abstract

Apical cell-cell junctions, including adherens junctions (AJs) and tight junctions (TJs), adhere epithelial cells to one another and regulate selective permeability at both bicellular junctions (BCJs) and tricellular junctions (TCJs). Although several specialized proteins are known to localize at TCJs, it remains unclear how actomyosin-mediated tension transmission at TCJs contributes to the maintenance of junction integrity and barrier function at these sites. Here, utilizing gastrula-stage *Xenopus laevis* embryos as a model system, we describe a mechanism by which Vinculin, a mechanosensitive protein, anchors the actomyosin network at TCJs, thus maintaining TJ stability and barrier function. Using an optogenetic approach, we found that acutely increasing junctional tension results in robust recruitment of Vinculin to apical junctions immediately surrounding TCJs. In Vinculin knockdown (KD) embryos, junctional actomyosin intensity is decreased and becomes disorganized at TCJs. Using fluorescence recovery after photobleaching (FRAP), we show that loss of Vinculin results in reduced Actin stability at TCJs. Vinculin knockdown also destabilizes Angulin-1, a key protein involved in regulating barrier function at TCJs. When Vinculin KD embryos are subjected to increased tension, TCJs cannot maintain their proper morphology. Finally, using a live imaging barrier assay, we detect increased barrier leaks at TCJs in Vinculin KD embryos. Together, our findings show that Vinculin-mediated actomyosin organization is required to maintain junction integrity and barrier function at TCJs and reveal new information about the interplay between adhesion and barrier function at TCJs.

**Highlights:**

- Vinculin is mechanosensitively recruited to tricellular junctions
- Vinculin’s actin-binding function is needed for tricellular actomyosin organization
- Tricellular tight junctions are unstable when Vinculin is knocked down
- Vinculin is required to maintain barrier function at tricellular junctions

## Introduction

Epithelial barrier function is essential for proper development, organ compartmentalization, and separation of internal and external environments [1]. Disruption of epithelial barriers can lead to pathogen invasion or diseases such as inflammatory bowel disease [2, 3]. Epithelia are made up of polarized epithelial cells connected by cell-cell junctions, including tight junctions (TJs) and adherens junctions (AJs) [2, 3]. TJs seal the paracellular space between cells and regulate the selective permeability characteristics of the tissue; AJs physically connect neighboring cells and mechanically link the cells within the tissue. AJs couple the cytoskeletons of neighboring cells by linking transmembrane cadherins to filamentous (F-) actin through catenins and other cytoplasmic linkers [4, 5]. One of these cytoplasmic linkers is the mechanosensitive protein, Vinculin, which binds both α-catenin and F-actin. At AJs experiencing high tension, α-catenin undergoes a conformational change, which reveals a binding site for Vinculin; Vinculin recruitment reinforces the AJ’s connection to F-actin [4, 6, 7]. Many studies have focused on Vinculin’s role in junction reinforcement at simple interfaces where only two cells touch (bicellular junctions, BCJs). In contrast, much less is known about how more complex multicellular junctions are maintained or reinforced when mechanical force is applied on epithelial tissues.

Tricellular junctions (TCJs), the points where three cells meet, are naturally sites of increased tension due to the tensile forces generated by the three BCJs and their associated F-actin and myosin (actomyosin) networks converging at a single TCJ [8, 9]. Despite this mechanical challenge on cell vertices, the cellular connections at these sites must be strong enough to maintain cell adhesion and barrier function under baseline tension and when mechanical force is applied on the tissue during developmental morphogenesis or tissue homeostasis [10].

Recent studies have identified unique molecular players known to localize specifically to TCJs. The first TCJ-specific protein identified was Gliotactin in the *Drosophila* epithelium [11]. Since then, additional TCJ-specific proteins have been identified in *Drosophila*: Sidekick at tricellular AJs and Anakonda and M6 at tricellular septate junctions (septate junctions in invertebrates are functionally analogous to TJs in vertebrates) [9, 12, 13]. The existence of these TCJ-specific proteins suggests that there may be unique mechanisms responsible for maintaining cell-cell connections and barrier function at TCJs. However, the junctional ultrastructure is different in invertebrates compared with vertebrates, and many of the TCJ-specific proteins identified in *Drosophila* (Gliotactin, Anakonda, M6) do not have vertebrate homologs [9, 12].

Within the vertebrate epithelium, two tricellular tight junction (tTJ) proteins have been identified that localize specifically to TCJs: angulins and Tricellulin. Angulin family proteins, including Angulin-1/LSR (lipolysis-stimulated lipoprotein receptor), localize to tTJs and recruit Tricellulin to tTJs [14, 15]. Both Angulin-1 and Tricellulin have been implicated in maintaining barrier function [12, 14, 15]. A recent study revealed a molecular connection between TJ and AJ complexes at TCJs and suggested that AJ proteins might play a role in regulating barrier function at TCJs [16]. The authors found that Tricellulin interacts directly with α-catenin, thus connecting TCJs to the actin cytoskeleton and supporting barrier function at TCJs [16]. However, many questions remain regarding the molecular players and mechanisms regulating vertebrate TCJs.

Vinculin is another likely candidate for facilitating the interplay between AJs and TJs at TCJs. Vinculin is concentrated at TCJs in *Xenopus* embryos, and its recruitment to TCJs is enriched when mechanical force is increased [9, 17]. Vinculin is also concentrated at TCJs in cultured Eph4 epithelial cells, while Vinculin knockout Eph4 cells exhibit a disrupted paracellular barrier for ions as well as “distorted” vertices [18]. Because Vinculin can reinforce the connection of AJs to F-actin in a force-dependent manner, we hypothesized that Vinculin might mechanosensitively strengthen TCJs.

In this study, we set out to test whether Vinculin regulates actomyosin-mediated tension transmission at TCJs and to determine Vinculin’s role in maintaining barrier function at these vulnerable sites in epithelia. Using gastrula-stage *Xenopus laevis* embryos as a model for the vertebrate epithelium, we validated a Vinculin morpholino to knock down Vinculin protein expression and added back either wildtype (WT) Vinculin or an actin-binding mutant of Vinculin. Using approaches including high-resolution microscopy, optogentically-activated mechanical force application, fluorescence recovery after photobleaching (FRAP) to measure protein stability at TCJs, and a live imaging barrier assay, we show that mechanosensitive recruitment of Vinculin to TCJs is needed for actomyosin organization and proper stability of the tTJ protein Angulin-1 at TCJs. Additionally, loss of Vinculin disrupts TCJ morphology and leads to barrier leaks specifically at TCJs. This work provides new insight into Vinculin’s role in maintaining junction integrity and barrier function at TCJs and adds valuable information about the interplay between cell-cell adhesion and barrier function.

## Results

### Vinculin is mechanosensitively recruited to TCJs

Vinculin is mechanosensitively recruited to focal adhesions [19, 20], AJs [4], the cleavage furrow of dividing cells [17], and TCJs [9, 17]. To better characterize and quantify Vinculin’s mechanosensitive recruitment to TCJs when mechanical force is applied, we used two complementary approaches to increase tension in gastrula-stage (Nieuwkoop and Faber stage 10-11 [21]) *Xenopus laevis* embryos: optogenetic activation of RhoA and addition of extracellular adenosine triphosphate (ATP) (**Fig 1A-C**).

**Figure 1:**
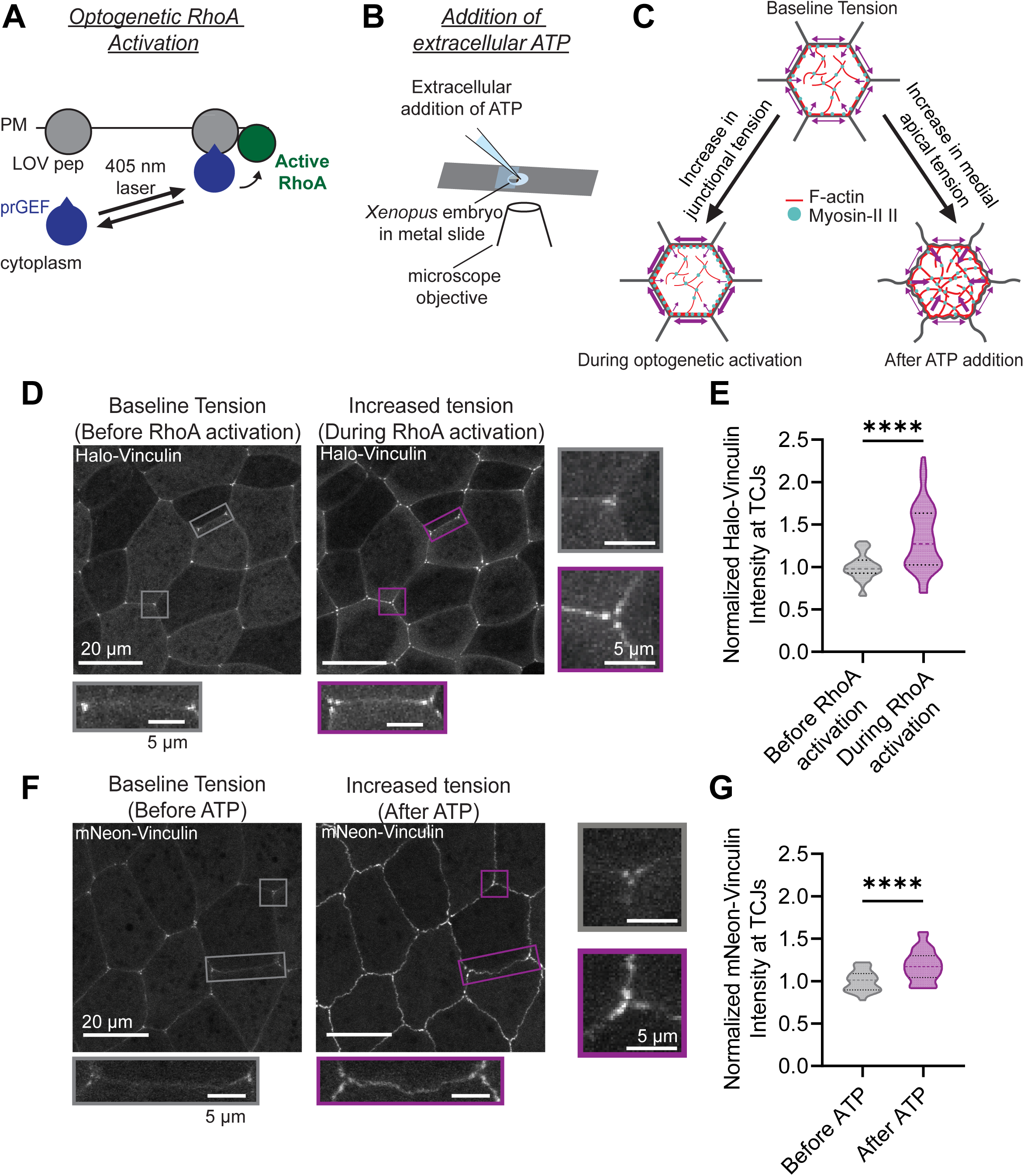
Vinculin is mechanosensitively recruited to TCJs. **A)** Schematic of the TULIP optogenetic system to activate RhoA. LOVpep is bound to the plasma membrane (PM). Upon 405 nm light stimulation, LOVpep undergoes a conformational change allowing it to interact with the photo-recruitable GEF (prGEF) and activate RhoA. **B)** Schematic of extracellular ATP addition. *Xenopus* embryos are mounted on a custom-made metal slide sandwiched between two glass coverslips. The top coverslip only covers 50% of the hole in the slide, allowing an opening to add ATP to the embryo during live confocal imaging. **C)** Schematic of observed cellular responses of optogenetic activation of RhoA and additional of extracellular ATP. Arrows represent expected forces, with thicker arrows representing more force. **D)** Live confocal images of epithelial cells expressing Vinculin (Halo-Vinculin with either JF646 or JFX 646), photo-recruitable GEF (prGEF-YFP), and LOVpep (GFP-silent-LOVpep). Images were captured before and during RhoA activation using optogenetic stimulation. Zoomed in panels (indicated by boxes) highlight changes in Vinculin recruitment at TCJs (right) and BCJs (bottom). **E)** Quantification of Halo-Vinculin intensity at TCJs before and during RhoA activation. Statistics, paired t-test; n = 3 experiments, 9 embryos, 55 TCJs; p ≤ 0.0001 (****). Violin plots show the median (dashed line) and the 25^th^ and 75^th^ quartiles (dotted lines). **F)** Live confocal images of cells expressing Vinculin (mNeon-Vinculin) before and after extracellular ATP addition. Zoomed in panels (indicated by boxes) highlight changes in Vinculin recruitment at TCJs (right) and BCJs (bottom). **G)** Quantification of mNeon-Vinculin intensity at TCJs before and after ATP addition. Statistics, paired t-test; n = 3 experiments, 5 embryos, 29 TCJs; p ≤ 0.0001 (****). Violin plots show the median (dashed line) and the 25^th^ and 75^th^ quartiles (dotted lines).

First, we used the TULIP optogenetic system [22, 23] that was previously adapted for use in *Xenopus* embryos [24] to increase contractility on demand. The TULIP system utilizes a photosensitive LOVpep domain that is anchored to the plasma membrane and a photo-recruitable guanine nucleotide exchange factor (prGEF) that activates RhoA upon stimulation with 405 nm light by recruiting the prGEF to the plasma membrane (**Fig 1A**). Active RhoA then activates downstream effectors resulting in increased junctional actomyosin contraction (**Fig S1A and A’**). Optogenetic activation of RhoA resulted in cell-cell junctions becoming more taut, consistent with increased tension along the junction (**Fig 1C and Video S1**). Tagged Vinculin was present weakly at BCJs and enriched at TCJs at baseline tension (**Fig 1D**). Following optogenetic RhoA activation, Vinculin was recruited to TCJs (33.6% increase in intensity) (**Fig 1D, E, and Video S1**), which was similar to the increase at BCJs (32.0% increase in intensity) (**Fig 1D, S2A, and Video S1**). This reveals that Vinculin’s recruitment to TCJs is mechanosensitive. Tagged Vinculin localized in three spots surrounding the TCJ (**Fig 1D**), as reported previously [9], where we proposed Vinculin helps anchor actomyosin bundles adjacent to TCJs. Upon optogenetic stimulation, the three Vinculin spots both increase in intensity and move away from the vertex (**Fig 1D and Video S1**).

As a complementary approach, we increased tension by adding extracellular ATP to *Xenopus* embryos while live imaging (**Fig 1B**), an approach which has previously been shown to increase contractility in *Xenopus* embryos [17, 25-27] Following the addition of extracellular ATP, Vinculin was mechanosensitively recruited to TCJs (18.1% increase in intensity) (**Fig 1F, G, and Video S2**). Extracellular ATP addition resulted in the relocalization of F-actin and Myosin II from the junctions to the medial-apical cortex (**Fig S1B and B’**). This relocalization results in wavy junctions, likely due to asymmetries in pulling forces on the BCJs generated by the medial-apical cortices in neighboring epithelial cells (**Fig 1C and Video S2**). Indeed, Vinculin was strongly recruited to BCJs upon addition of extracellular ATP (38.9% increase in intensity) (**Fig 1F, S2B, and Video S2**). Prior to ATP addition, Vinculin again localized in three spots around TCJs, which increased in intensity and became elongated upon ATP addition. In contrast to Vinculin’s localization, the tTJ protein Angulin-1 forms a single spot at the vertex and does not separate around the TCJ with the addition of extracellular ATP (**Fig 1B”**). This suggests that the tTJ appears intact as Vinculin is mechanosensitively recruited around TCJs.

Interestingly, there was a larger increase in mechanosensitive recruitment of Vinculin to TCJs in response to optogenetic stimulation than in response to extracellular ATP addition (33.6% vs. 18.1% increase, respectively). This data suggests that tension distribution differs between these two approaches for increasing contractility. Together, this data indicates that Vinculin is recruited to TCJs in a tension-dependent manner.

### Vinculin actin-binding mutant disrupts TCJ actomyosin organization

In order to investigate Vinculin’s functional role at TCJs in *Xenopus* embryos, we knocked down Vinculin using a custom-designed antisense morpholino oligomer that binds to the 5’ untranslated region (UTR) of Vinculin mRNA and blocks translation of endogenous Vinculin (**Fig S3A**). Immunofluorescence with an anti-Vinculin antibody validated effective Vinculin knockdown (KD), which could be rescued by injection of wildtype (WT) or mutant Vinculin mRNAs that could not be targeted by the morpholino (**Fig S3C and D**). Additionally, Vinculin KD resulted in increased cell surface area, and exogenous expression of WT Vinculin expression largely rescued this effect (**Fig S3E**).

Next, we tested how the loss of Vinculin, or its actin-binding function, affected actomyosin organization at TCJs. In control embryos, F-actin was concentrated in an apical bundle encircling each epithelial cell, whereas Myosin II was localized in a “train track” pattern on either side of the junction (**Fig 2A**). Upon Vinculin KD, the intensity of F-actin and Myosin II at TCJs was significantly decreased (**Fig 2A-C**) but could be rescued when WT Vinculin mRNA was injected into Vinculin KD embryos (**Fig 2A-C**). To investigate the role of Vinculin’s actin-binding function in maintaining TCJ cytoskeletal organization, we utilized R1049E, a point mutation in Vinculin that has been shown to decrease actin binding by 6-fold without affecting Vinculin’s structure [28]. Vinculin is highly conserved between species (the mouse sequence is 99% similar to human, while the frog sequence is 96% similar to human), and this actin-binding region, including residue R1049, is 100% conserved between humans, mice, and frogs (**Fig S3B**). Vinculin R1049E localized to cell-cell junctions (**Fig S3C and D**), suggesting that the mutation does not impair Vinculin’s recruitment to cell-cell junctions in frog embryos. When Vinculin R1049E mRNA was injected into Vinculin KD embryos, F-actin and Myosin II intensity at TCJs was reduced compared with WT Vinculin rescue (**Fig 2A-C and S3C**). Interestingly the Myosin II “train track” patterning is lost in Vinculin KD embryos and rescued in KD+WT embryos (**Fig 2D**). However, KD+R1049E results in a partial rescue where Myosin II is localized to TCJs, but not organized correctly, with the line scan showing a single peak as opposed to the two peaks observed in the controls (**Fig 2D**). Together, this data suggests that Vinculin organizes actomyosin at TCJs via its direct interaction with actin filaments.

**Figure 2:**
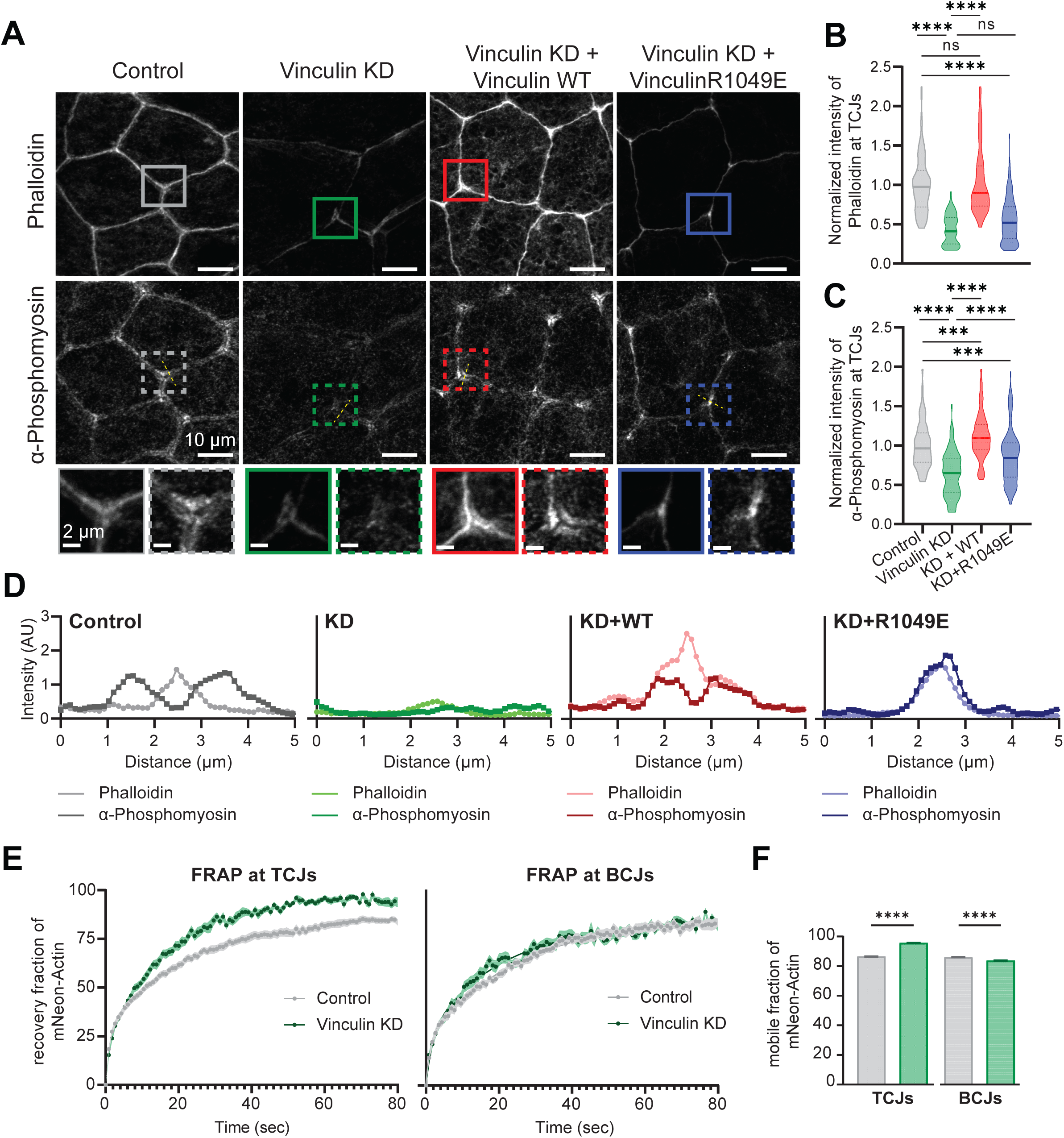
Vinculin actin-binding mutant disrupts TCJ actomyosin organization and stability. **A)** Fixed confocal images of epithelial cells from control embryos, Vinculin knockdown embryos (Vinculin KD), Vinculin knockdown embryos rescued with wildtype Vinculin mRNA (KD + WT), and Vinculin knockdown embryos rescued with mRNA encoding an actin-binding mutant of Vinculin (KD + R1049E) that were stained for F-actin (phalloidin Alexa Fluor 568) and phosphomyosin (α- phosphomyosin light chain 2 antibody). Zoomed in panels (indicated by boxes) highlight changes at TCJs. **B)** and **C)** Quantification of phalloidin and α-phosphomyosin intensity at TCJs. Statistics, one-way ANOVA; n = 3 experiments; control = 174 TCJs, 25 embryos; Vinculin KD = 90 TCJs, 14 embryos; KD + WT = 161 TCJs, 23 embryos; KD + R1049E = 140 TCJs, 20 embryos; p-values > 0.05 (ns), ≤ 0.001 (***), ≤ 0.0001 (****). Violin plots show the median (solid line) and the 25^th^ and 75^th^ quartiles (dotted lines). **D)** Line scans of phalloidin and α-phosphomyosin adjacent to TCJs in control, Vinculin KD, KD + WT, and KD + R1049E embryos. Locations of line scans are indicated by dashed yellow lines in (**A**). **E)** *Left:* Recovery curve (dots) from mNeon-Actin FRAP at TCJs with a double exponential nonlinear fit (solid line). n = 3 experiments; control = 8 embryos, 21 TCJs; Vinculin KD = 7 embryos, 18 TCJs; errors bars, SEM. *Right:* Recovery curve (dots) from mNeon-Actin FRAP at BCJs with a double exponential nonlinear fit (solid line). n = 3 experiments; control = 7 embryos, 20 BCJs; Vinculin KD = 8 embryos, 23 BCJs. **F)** Mobile fractions calculated from (**E**). Statistics, unpaired t-test; p ≤ 0.0001 (****); error bars, SEM.

### Vinculin stabilizes actin at TCJs

We next asked whether Vinculin affects actin stability at TCJs and/or BCJs. To answer this question, we compared FRAP curves of actin (mNeon-Actin) in control and Vinculin KD embryos (**Fig 2E**). From the FRAP curves, we calculated the mobile fraction, a measure of *how much* actin turns over, and t_1/2_, a measure of *how fast* actin turns over, at both TCJ and BCJ locations (**Fig S4A**). At TCJs, the mobile fraction for actin was significantly increased in Vinculin KD embryos compared with control embryos (95.7% vs. 86.8%, respectively; **Fig 2F and S4B**), indicating that actin is stabilized at TCJs in the presence of Vinculin. FRAP data was fitted with a double exponential curve to derive the fast and slow halftimes of recovery (t_1/2_). The slow t_1/2_ was significantly smaller in Vinculin KD embryos compared to control embryos (12.3s vs. 16.9s, respectively; **Fig S4B**), and the fast t_1/2_ followed a similar trend (0.69s vs. 0.74s, respectively; **Fig S4B**), indicating that mNeon-Actin signal recovers faster when Vinculin is knocked down. In contrast to TCJs where the mobile fraction was increased by 10.6% when Vinculin was KD, at BCJs, the mobile fraction was actually decreased by 2.67%, and the fast and slow t_1/2_ were not significantly changed (**Fig 2E, 2F and S4B**). These FRAP results indicate that loss of Vinculin results in a more dynamic pool of actin specifically at TCJs. Combined with the fixed staining data for actomyosin organization, our results suggest Vinculin regulates proper actomyosin architecture and actin stability at TCJs.

### Vinculin is required for maintaining tricellular tight junction protein stability

We next investigated how Vinculin-mediated changes in actomyosin organization and actin stability affect tTJ integrity. Angulin-1 is a critical tTJ component and is essential for maintaining barrier function at TCJs [29, 30]. Since Angulin-1 is the core protein at the tTJ and recruits Tricellulin [9, 12], we used Angulin-1 as a representative tTJ protein. Using immunofluorescence, we found that Angulin-1 intensity at TCJs was slightly but significantly decreased in Vinculin KD embryos compared to control embryos (**Fig 3A and 3B**), although the overall protein expression level was not detectably different in Vinculin KD embryos and controls (**Fig S3F**). FRAP of Angulin-1 (Angulin-1-3xGFP) at TCJs revealed that Angulin-1 recovered faster and the mobile fraction was significantly higher in Vinculin KD embryos compared to control embryos (mobile fraction: 63.1% vs. 49.4%, respectively; **Fig 3C-E and S4C and Video S3**). Both the slow t_1/2_ and fast t_1/2_ were significantly smaller in Vinculin KD embryos compared to control embryos (slow t_1/2_: 38.1s vs. 83.8s; respectively; **Fig 3F and S4C**; fast t_1/2_: 1.43s vs. 4.72s, respectively; **Fig 3G and S4C**). Together, this FRAP data indicates that Vinculin stabilizes Angulin-1 at tTJs.

**Figure 3:**
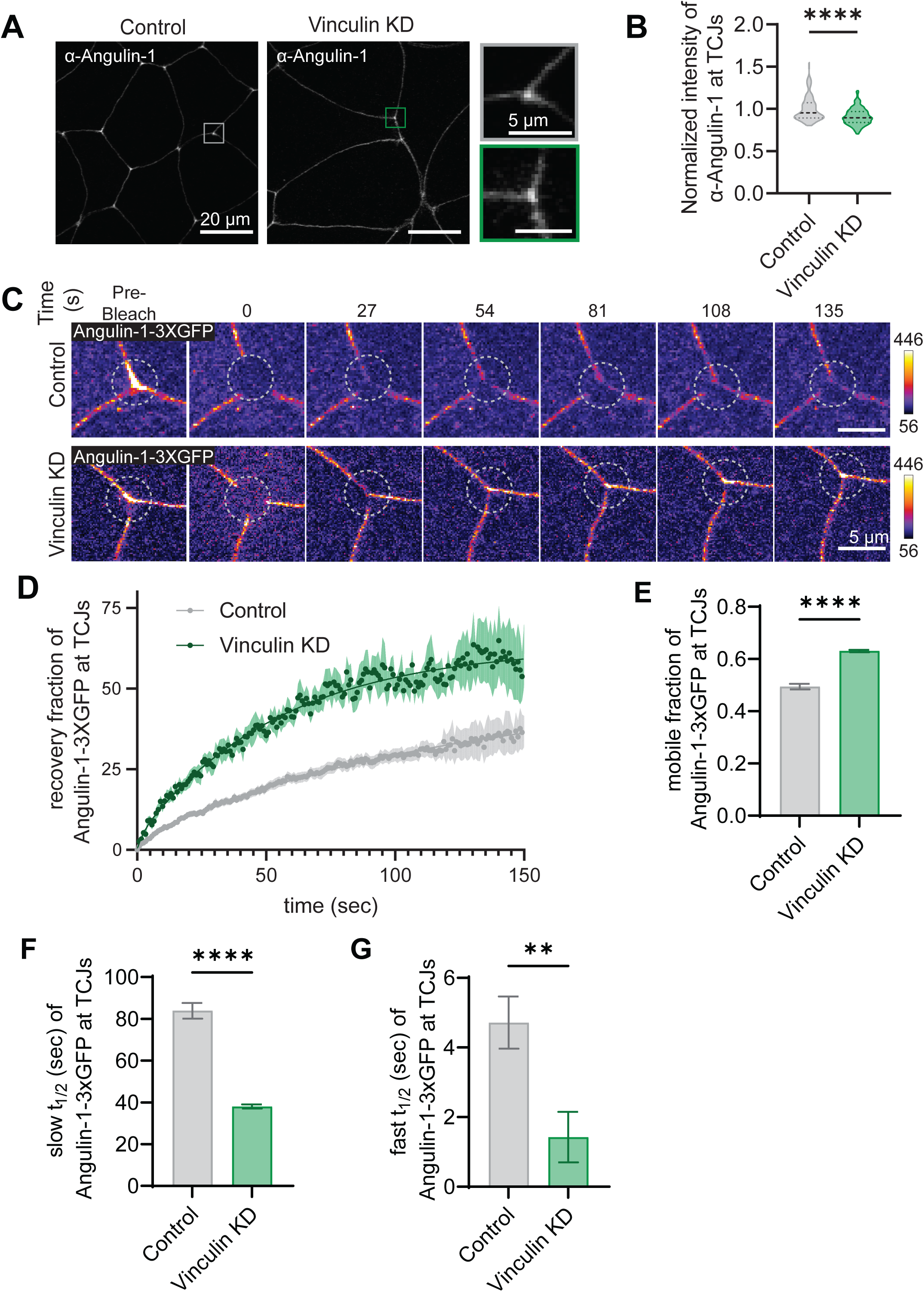
Vinculin is required for maintaining tricellular tight junction protein stability. **A)** Fixed confocal images from control and Vinculin KD embryos that were stained for Angulin-1 (α-Angulin-1). **B)** Quantification of the intensity α-Angulin-1 at TCJs. Statistics, unpaired t-test with Welch’s correction; n = 3 experiments; control = 23 embryos, 170 TCJs; Vinculin KD = 25 embryos, 185 TCJs; p ≤ 0.0001 (****). Violin plots show the median (dashed line) and the 25^th^ and 75^th^ quartiles (dotted lines). **C)** Montage of Angulin-3xGFP FRAP in control and Vinculin KD embryos pre-bleach and at times 0, 27, 54, 81, 108, 135 seconds after bleaching. Dashed circle indicates photobleached region. Images are shown using the FIRE lookup table (LUT). LUTs were adjusted in the same way for each image. **D)** Recovery curve (dots) from Angulin-3xGFP FRAP at TCJs with a double exponential nonlinear fit (solid line). n = 3 experiments: control = 8 embryos, 15 TCJs; Vinculin KD = 6 embryos, 9 TCJs; errors bars, SEM. **E)** Mobile fraction calculated from (**D**). Statistics, unpaired t-test with Welch’s correction; p ≤ 0.0001 (****); error bars, SEM. **F)** Slow t_1/2_ calculated from (**D**). Statistics, unpaired t-test with Welch’s correction; p ≤ 0.0001 (****); error bars, SEM. **G)** Fast t_1/2_ calculated from (**D**). Statistics, unpaired t-test with Welch’s correction; p ≤ 0.01 (**); error bars, SEM.

### Vinculin maintains TCJ morphology under increased tension

Because we observed disrupted actomyosin organization at TCJs along with disrupted tTJ protein stability at baseline tension when Vinculin is knocked down, we hypothesized that Vinculin KD embryos would be more susceptible to mechanical failure at TCJs when stressed. To further mechanically stress TCJs, we optogenetically activated RhoA as described earlier (**Fig 1A**). Under increased tension, TCJs in control embryos remained unchanged in morphology before and during RhoA activation. However, there was a significant increase in “dipped” TCJs when Vinculin was knocked down (**Fig 4A-D, Video S4**). When confocal images are Z-projected (Full Z-Projection), this difference is difficult to appreciate. However, these differences become evident when viewing Z-projections of the most apical slices (Apical Z-Projection), where the dipped TCJs appear as “holes” at the TCJs (**Fig 4A**). The increase in “dipped” TCJs when tension is applied on Vinculin KD embryos is also apparent when viewing the XZ view of a TCJ (**Fig 4A**) or a 3D projection of the tissue (**Fig 4B**). We then quantified the distance TCJs dipped in controls compared to Vinculin KD embryos (**Fig 4C**). Control TCJs exhibited a slight but not significant difference in the distance TCJs dip before and during RhoA activation (0.37 μm to 0.90 μm on average). In contrast, Vinculin KD TCJs significantly differed in the distance TCJs dip before and during RhoA activation (2.78 μm to 5.04 μm on average) (**Fig 4D**). Together, this data indicates that TCJ morphology is severely disrupted in Vinculin KD embryos and that this is further exasperated when under increased tension. Along with our earlier finding that Vinculin KD results in a significant decrease in actomyosin at TCJs, these results suggest that the actomyosin architecture that Vinculin maintains at TCJs is essential for preventing TCJs from deforming under increased tension.

**Figure 4:**
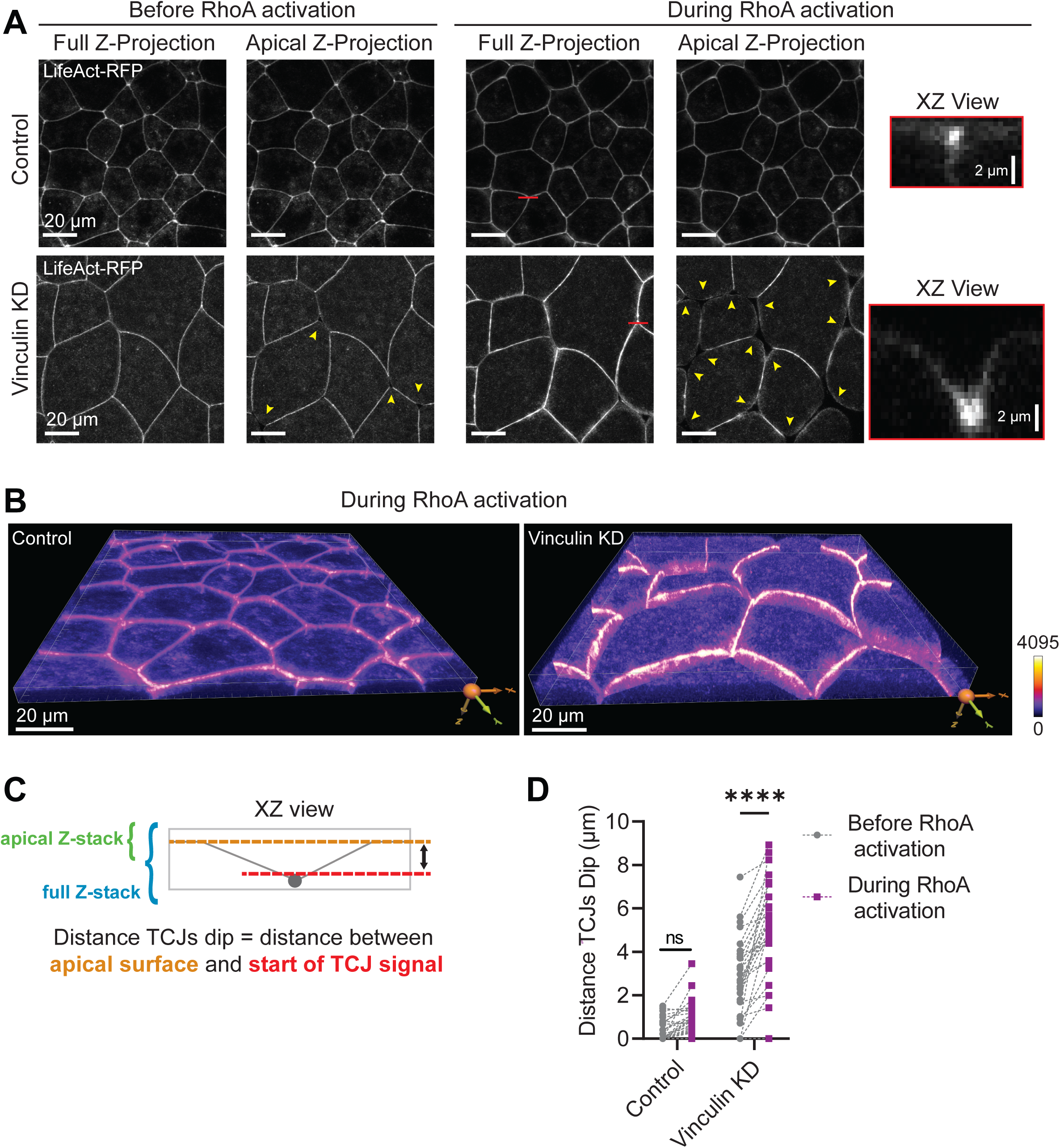
Vinculin is required for maintaining TCJ morphology. **A)** Live confocal images of control and Vinculin KD embryos that express an F-actin probe (LifeAct-RFP), photo-recruitable GEF (prGEF-YFP), and LOVpep (GFP-silent-LOVpep). Images of full Z-projections and apical Z-projections are shown at both baseline (before RhoA activation) and increased Rho-mediated tension (during RhoA activation). Yellow arrowheads point to dipped TCJs. Red bars indicate the TCJs that are shown in XY views on the right in red boxes. **B)** 3D views of control and Vinculin KD embryos during RhoA activation from (**A**). Images are shown using the FIRE lookup table (LUT). LUTs were adjusted in the same way for each image. **C)** Schematic highlighting the difference between “apical Z-stack” and “full Z-stack” as well as how distance TCJs dip was calculated. **D)** Quantification of distance TCJs dip before and during RhoA activation in both control and Vinculin KD embryos. Statistics, paired t-test; n = 3 experiments; control = 4 embryos, 28 TCJs; Vinculin KD = 4 embryos, 28 TCJs; p-values > 0.05 (ns), ≤ 0.0001 (****).

### Vinculin is required for maintaining barrier function at TCJs

Based on our observation that in Vinculin KD embryos, Angulin-1 is less stable at tTJs, and TCJs under increased tension appear deformed, we next wanted to test if loss of Vinculin resulted in defects in epithelial barrier function – particularly at TCJs. To measure local changes in barrier function, we used the Zinc-based Ultrasensitive Microscopic Barrier Assay (ZnUMBA) [31, 32]. This assay allows measurement of localized barrier leaks with high spatiotemporal resolution (**Fig 5A**). A fluorogenic Zn^2+^ indicator, FluoZin3, is injected into the blastocoel of gastrula-stage embryos, and embryos are bathed in media containing Zn^2+^. If the epithelial barrier is breached, FluoZin3 and Zn^2+^ can interact, resulting in an increase in FluoZin3 fluorescence intensity. Notably, Vinculin KD embryos frequently exhibited high FluoZin3 signal at TCJs compared with controls (**Fig 5B and Video S5**), indicating that Vinculin KD embryos have impaired barrier function specifically at TCJs. By measuring the percentage of TCJs that leaked during 25-minute videos, we found that Vinculin KD embryos had a significantly higher percentage of leaky TCJs compared to control embryos (51.6% ± 10.0% vs. 10.3% ± 3.2%, respectively; **Fig 5C**). The leaks at TCJs in Vinculin KD embryos generally appeared and were resolved over the course of a 25- minute movie, indicating that these leaks were transient (**Fig 5D and Video S5)**. Together with our findings that Vinculin regulates actomyosin architecture at TCJs and stabilizes Angulin-1 at tTJs, this data suggests that Vinculin is important for maintaining junction integrity and epithelial barrier function at TCJs.

**Figure 5:**
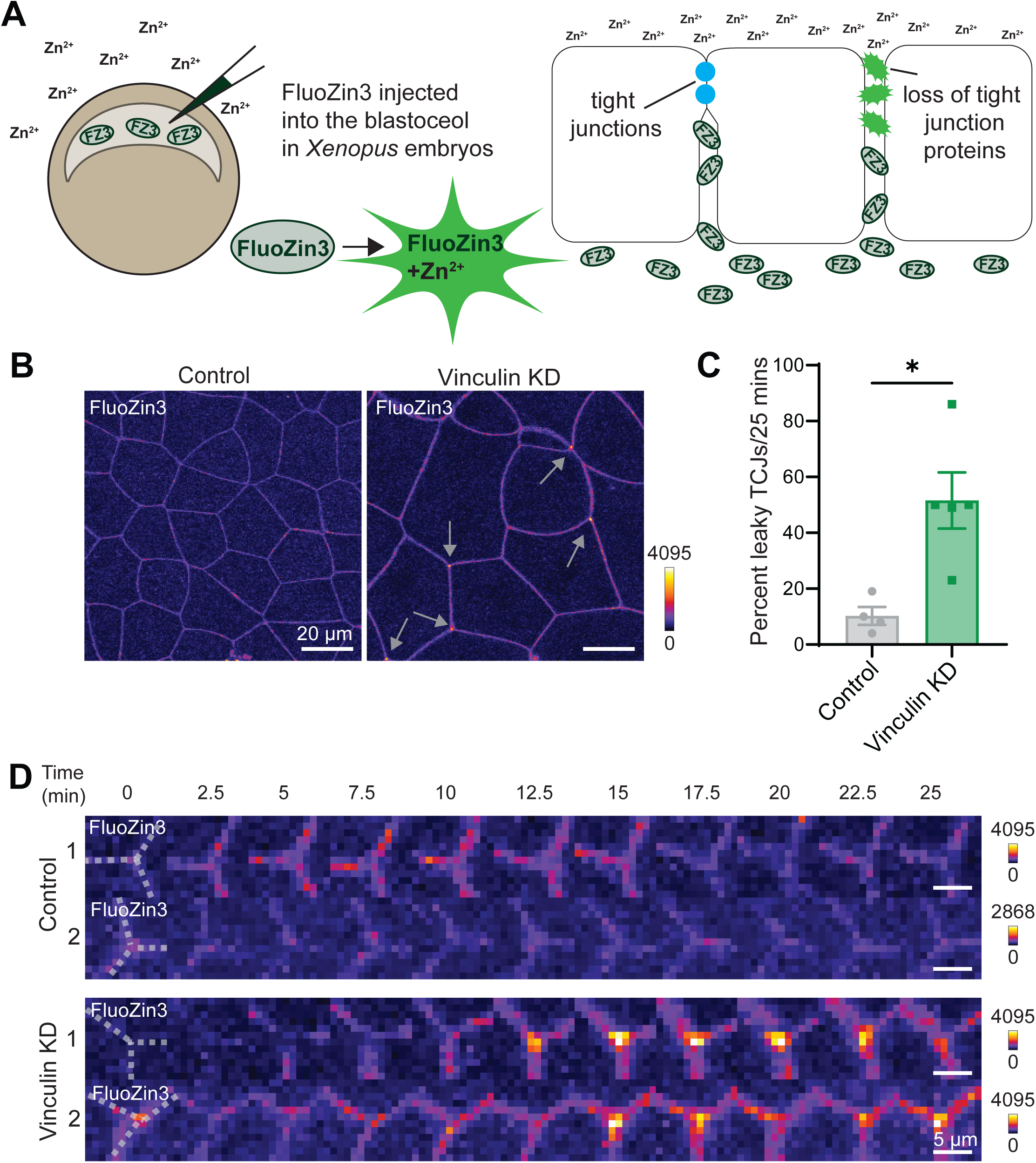
Vinculin is required to maintain barrier function at TCJs. **A)** Schematic of Zinc-base Ultrasensitive Microscopic Barrier Assay (ZnUMBA). Disruption of tight junction proteins leads to interaction between Zn^2+^ and FluoZin3, causing increased FluoZin3 fluorescence and indicating a leak in barrier function. **B)** Live confocal images of FluoZin3 signal in control and Vinculin KD embryos. Images are shown using the FIRE LUT, adjusted in the same way for each image. Gray arrows point to increased FluoZin3 signal at TCJs. **C)** Quantification of the percent of TCJs that showed an increase of FluoZin3 signal at TCJs during 25-minute movies. Images are shown using the FIRE LUT; adjustments are indicated for each image. Statistics, unpaired t-test with Welch’s correction; n = 3 experiments; control = 4 embryos, 223 TCJs; Vinculin KD = 5 embryos, 133 TCJs; p-value ≤ 0.05 (*); error bars, SEM. **D)** Montages of FluoZin3 signal at representative TCJs in control and Vinculin KD embryos. Images are shown using the FIRE LUT; adjustments are indicated for each image.

## Discussion

TCJs are naturally tension hotspots, and when epithelial tissues are mechanically challenged, TCJs become even more vulnerable to disruption. Despite their integral role in regulating tissue integrity and barrier function at the vertices where three cells meet, many unanswered questions remain regarding the molecular players and mechanisms regulating vertebrate TCJs. In this study, we used the intact embryonic epithelium of *Xenopus* embryos to better understand TCJ architecture and barrier function. Our work reveals that Vinculin plays a unique and critical role in maintaining TCJ integrity and barrier function specifically at TCJs via its direct interactions with F-actin. This is notable because aside from a recent study from our group that detected local leaks at TCJs when Angulin-1 was knocked out [31], previous studies only were able to identify global changes to barrier function when tTJ proteins Tricellulin [15, 16] or Angulin1 [14, 33, 34] were perturbed. Furthermore, our work suggests that the AJ protein Vinculin helps regulate the interplay between AJs and TJs at TCJs. We first show that Vinculin is mechanosensitively recruited to TCJs. Second, when Vinculin is knocked down, F-actin and Myosin II are reduced at TCJs, Myosin II is disorganized, and actin and Angulin-1 both exhibit reduced stability at TCJs. Third, TCJ morphology is disrupted in Vinculin KD embryos when challenged with additional tension. Finally, we show that Vinculin is required to maintain epithelial barrier function at TCJs. Thus, we define a mechanism in which junctional integrity and barrier function at TCJs require mechanosensitive recruitment of Vinculin, which mediates proper actomyosin organization at TCJs (**Fig 6A and B**).

**Figure 6:**
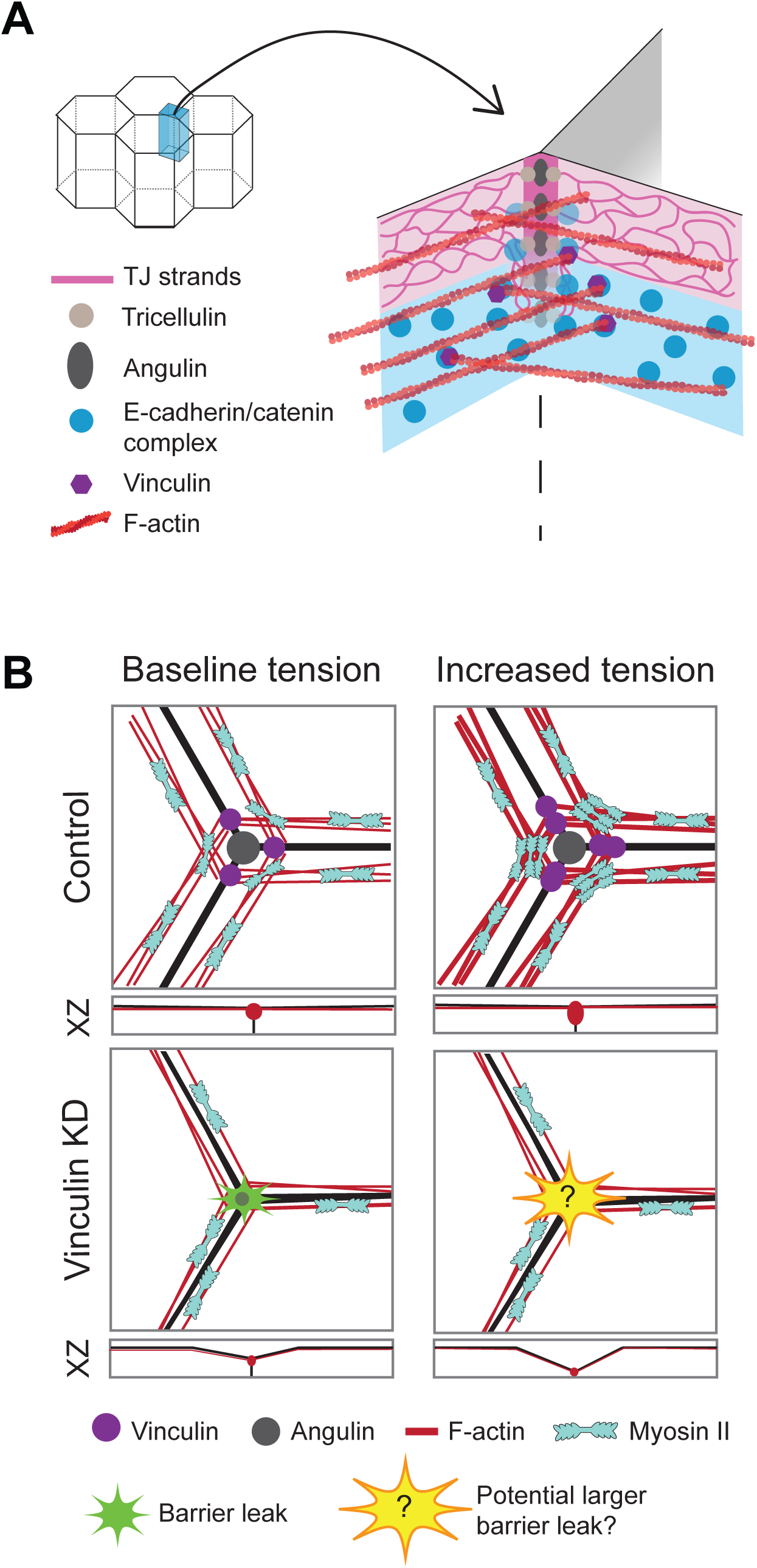
Model for Vinculin’s role at TCJs. **A)** 3D model of a TCJ showing Vinculin anchoring actin bundles at the TCJ to maintain junctional integrity. **B)** Top-down view of TCJs at baseline and increased tension for both controls and Vinculin KD embryos. Vinculin is mechanosensitively recruited to TCJs. When Vinculin is lost, actomyosin is disorganized, tricellular TJ proteins are less stable, and TCJs have an increased frequency of barrier leaks.

### Vinculin anchors actomyosin at TCJs

TCJs are sites of increased tension relative to BCJs; however, the mechanism of tension transmission between the junctional complex and the actin cytoskeleton at TCJs remains unclear. Several studies suggest that actomyosin bundles make end-on connections with cadherin-catenin complexes at TCJs [9, 16, 35, 36], distributing tension around the vertex. We previously suggested that Vinculin could be a critical linker connecting actomyosin to TCJs [17]. Our results here indicate that Vinculin anchors actomyosin bundles at TCJs, which is vital for proper tension transmission at TCJs. When tension is increased, more Vinculin is mechanosensitively recruited to strengthen the connection of actomyosin to the tricellular AJ. When Vinculin is knocked down, the actomyosin array at TCJs is disrupted and proper TCJ morphology is not maintained when tension is increased, suggesting that Vinculin-mediated connections to actomyosin are important for proper TCJ architecture and responsiveness to mechanical force. Recent single molecule studies demonstrated that under high tension, Vinculin’s association with the cadherin-catenin complex in the cytoplasm allosterically converts the interactions of the extracellular E-cadherin domains on opposing cells from weak “X-dimers” into strong “strand swap” dimers [37]. We speculate that the increased accumulation of Vinculin we have observed both in this study and previously [9, 17], along with others [18], could both anchor actomyosin to TCJs and allosterically strengthen E-cadherin-mediated adhesion at TCJs.

To further investigate Vinculin’s role in organizing actomyosin at TCJs, we tested whether the changes in actomyosin organization caused by Vinculin KD were due to Vinculin’s interaction with F-actin (as opposed to Vinculin’s interaction with other known actin-binding proteins). Injecting WT Vinculin mRNA rescued the actomyosin defects in Vinculin KD embryos; however, injecting a previously-characterized actin-binding mutant of Vinculin, R1049E [28], resulted in actomyosin disruptions comparable to when Vinculin is knocked down (**Fig 2**). These results further support the model that Vinculin directly anchors and organizes actomyosin at TCJs through its actin-binding capabilities (**Fig 6A and B**).

### Interplay between TJs and AJs

Our study reveals new data supporting the interplay between TJs and AJs. Traditionally, TJs and AJs have been studied as complexes that function independently from one another: TJs being responsible for barrier function and AJs being responsible for adhering and mechanically coupling cells. However, more recent studies have revealed an interdependency between the two complexes. Indeed, Vinculin has been shown to interact with the TJ protein ZO-1 [18, 38], allowing for a molecular connection between AJs and TJs. Another study found that the AJ protein α- catenin directly interacts with the tTJ protein Tricellulin, thus facilitating the connection between the tTJ and actomyosin [16]. While we propose a model where Vinculin maintains TCJ integrity and barrier function through actomyosin organization (**Fig 6**), it is also possible that Vinculin can regulate tTJs via direct interactions with ZO-1 and the Tricellulin-catenin complex.

Several recent studies have found that AJ proteins can directly impact barrier function. One study showed that Vinculin was essential for maintaining barrier function against ions in an epithelial cell line [18]. These authors found that barrier function defects in Vinculin knockout cells were rescued when tension was decreased using blebbistatin, suggesting that Vinculin normally helps resist mechanical forces that can disrupt barrier function. Other studies showed that Afadin, a scaffolding protein that links AJs and the actin cytoskeleton, can also interact with ZO-1 [39] and is necessary for maintaining barrier function under high tension [35]. Our findings reveal that Angulin-1 stability at TCJs is disrupted in Vinculin KD embryos. Additionally, when Vinculin is lost, barrier leaks specifically at TCJs are increased. These results support the idea that TJ and AJ functions are highly interdependent – especially at TCJs.

### Mechanotransduction at TCJs

Recent studies have shed new light on the dynamic properties of TCJs. It is becoming clear that TCJs can sense and respond to changes in mechanical forces during epithelial remodeling processes including morphogenesis, cell division, and tissue-level contraction. Our results show that Vinculin is mechanosensitively recruited to TCJs when actomyosin-mediated contractility is acutely increased. This is likely due to Vinculin binding to the mechanically stretched conformation of α-catenin [4]. Indeed, an antibody that binds specifically to mechanically stretched α-catenin is enriched at TCJs in cultured epithelial cells [18]. Similar to α-catenin, ZO-1 was recently shown to be mechanosensitive, changing to a stretched conformation in response to high tension [40]. This conformational change in ZO-1 mechanosensitively recruits Occludin and a transcription factor to TJs. Interestingly, double-knockdown of both ZO-1 and the closely related protein ZO-2 increases junctional and apical epithelial tension [35, 41], suggesting that ZO-1 normally helps mitigate mechanical stress in epithelial cells. Like Vinculin, Afadin plays a role in mechanosensitively strengthening the link between tricellular AJs and actomyosin. Afadin is strongly enriched at TCJs in ZO-1/ZO-2 double-knockdown cultured epithelial cells and is important for maintaining adhesion and actomyosin architecture at TCJs [35]. In *Drosophila*, Canoe (the *Drosophila* homolog of Afadin), plays an essential role in linking AJs to actin during the dynamic morphogenetic movements of development [35, 42]. A recent study showed that Canoe’s localization to TCJs is mechanosensitive and enhanced by Abl tyrosine kinase-mediated phosphorylation of Canoe [43]. This dynamic mechanosensitive localization of Canoe to cell vertices is necessary for proper cell rearrangements during cell intercalation [43].

TCJs can also sense and respond to reduced mechanical forces and adhesion. During *Drosophila* oogenesis, the follicular epithelium undergoes a process called patency, where TCJs remodel to allow for paracellular transport of yolk proteins through the epithelium for uptake by the oocyte [44, 45]. During this process, TCJs intentionally open transiently to allow for the transport of yolk proteins. A recent study showed that the opening of TCJs during patency is preceded by the sequential removal of several adhesion proteins and reduced actomyosin contractility [46]. Additionally, when the authors artificially stabilized AJs, they were able to prevent patency solely through modulating cell adhesion [46]. Related to our findings, this research highlights that dynamic regulation of adhesion and actomyosin contractility at TCJs is essential for regulating barrier function at TCJs.

## Conclusion

In this study, we demonstrate a novel role for Vinculin in maintaining epithelial integrity and barrier function at TCJs in *Xenopus laevis* gastrula-stage embryos by helping anchor actomyosin bundles at TCJs. Vinculin’s ability to be mechanosensitively recruited to TCJs under increased mechanical force may be especially important in dynamic epithelial tissues that need to tune their adhesion and maintain overall barrier function during developmental morphogenesis or in adult epithelial tissues that experience high mechanical forces.

## STAR Methods

### Key resources table

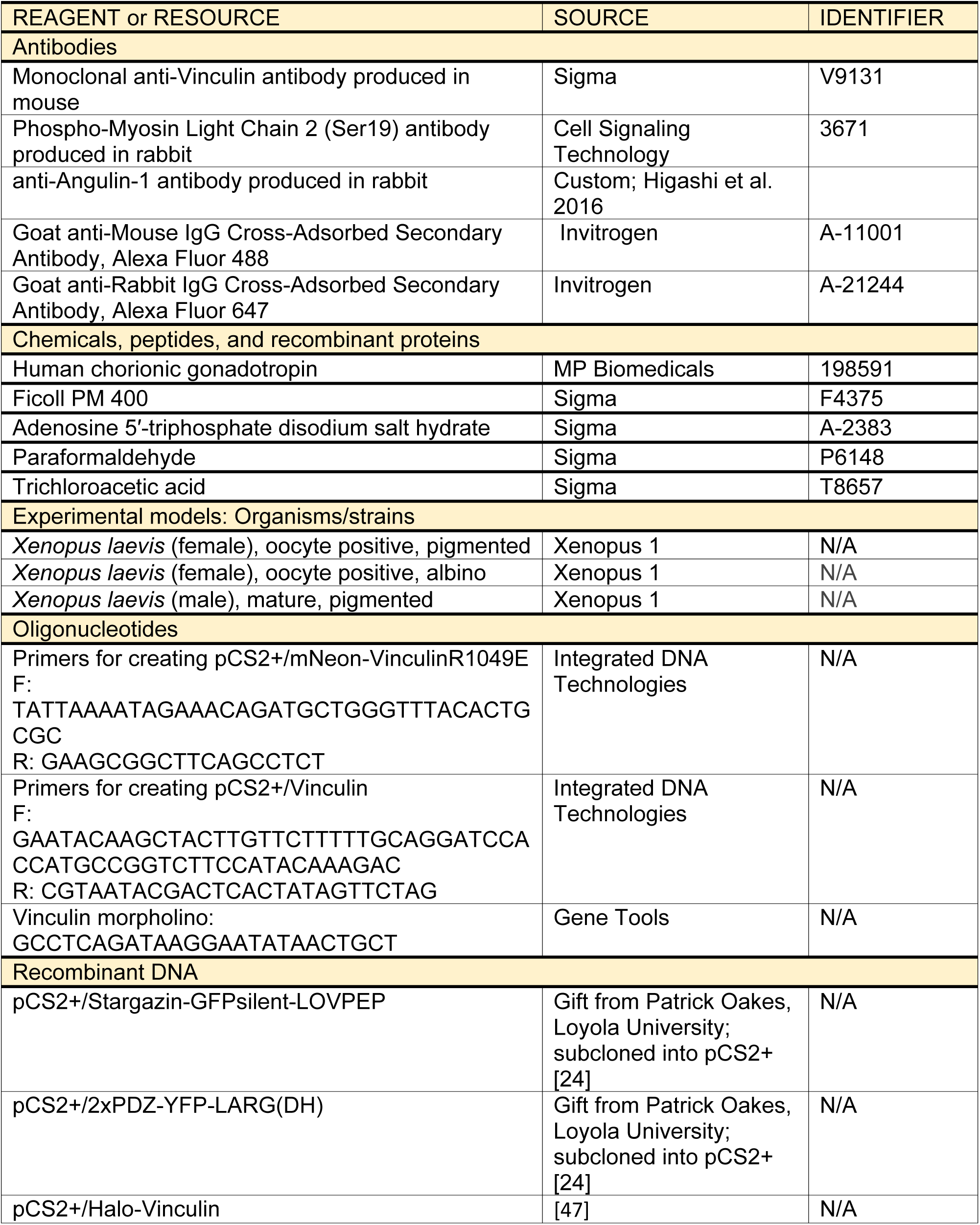
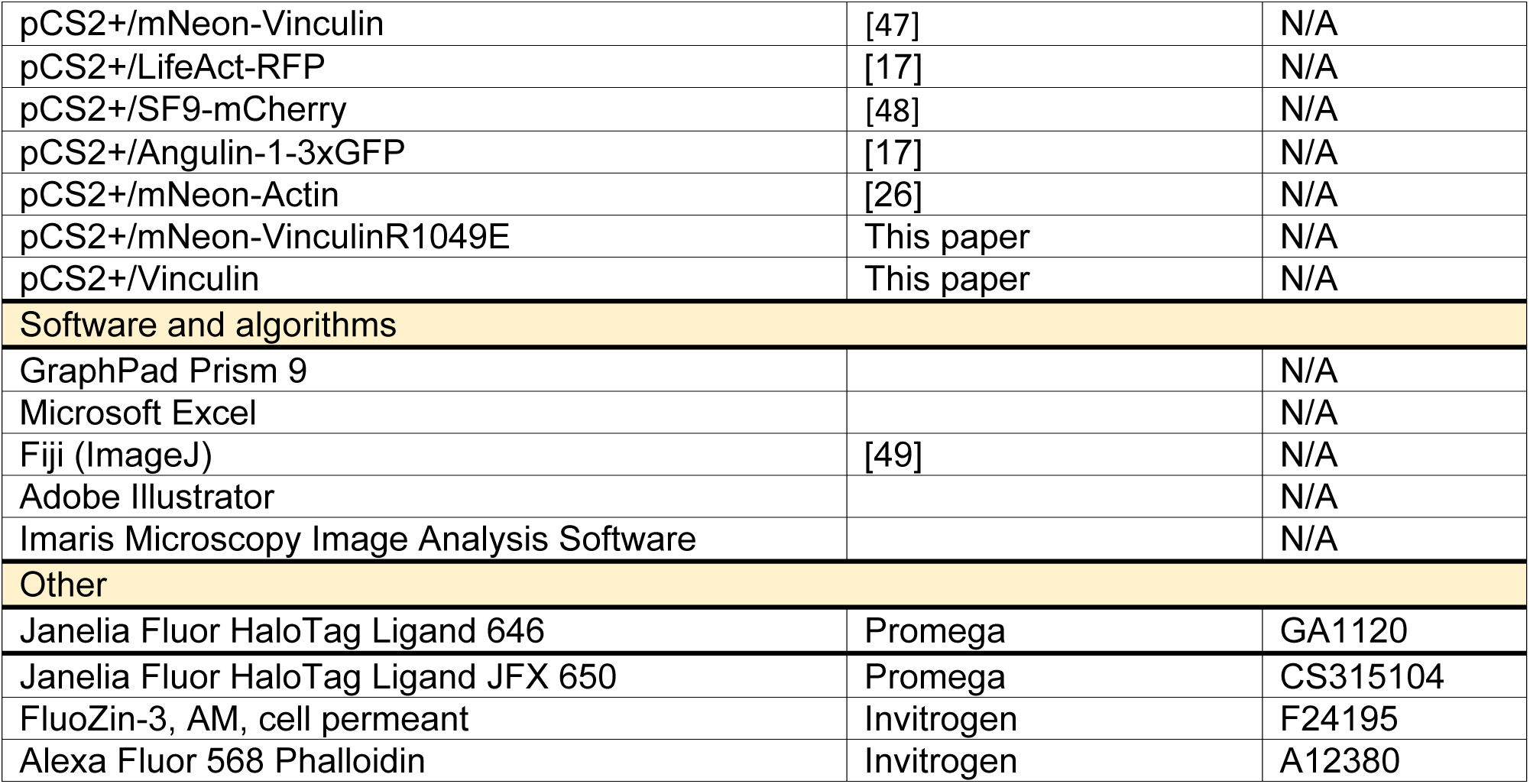

### Resource availability

#### Lead contact

Further information and requests for resources and reagents should be directed to and will be fulfilled by the Lead Contact, Ann L. Miller

#### Materials availability

Plasmids generated in this study can be obtained from the Lead Contact, Ann L. Miller

#### Data and code availability

All microscopy data reported in this paper will be shared by the lead contact upon request. This paper does not report original code. Any additional information required to reanalyze the data reported in this paper is available from the lead contact upon request.

#### Experimental model and subject details

Adult *Xenopus laevis* female frogs (wildtype and albino) and wildtype male frogs were purchased from Xenopus1 (Dexter, MI). The frogs were housed in a temperature-controlled aquatics facility in recirculating tank systems (Tecniplast, Milan, Italy), which maintain parameters for optimal water quality for the frogs (temperature, pH, and conductivity). Female frogs were injected with hormone (human chorionic gonadotropin) to induce them to lay eggs. Male frogs’ testes were harvested and used for egg fertilization.

After egg collection, *Xenopus* eggs were fertilized *in vitro*, dejellied in 2% cysteine, pH 7.8 in 1X Mark’s Modified Ringer’s solution (MMR) (5 mM HEPES, 100 mM NaCl, 2 mM KCl, 1 mM MgCl_2_, 2 mM CaCl_2_, pH 7.4), and stored in 0.1X MMR [50]. At 2-cell or 4-cell stage, the embryos were injected with mRNAs to express wildtype, mutant, or fluorescently-tagged proteins of interest or injected with morpholino (MO) to knock down Vinculin in 0.1X MMR or 0.1X MMR with 5% Ficoll. Embryos were kept in 0.1X MMR overnight at 15°C and then imaged at gastrula-stage (Nieuwkoop and Faber stage 10-11, [21]) by live or fixed microscopy techniques. Alternatively, gastrula-stage embryos were used to generate lysates for Western blotting.

All animal procedures strictly adhere to the compliance standards of the US Department of Health and Human Services Guide for the Care and Use of Laboratory Animals and were approved by the Institutional Animal Care and Use Committees at the University of Michigan. A board-certified Laboratory Veterinarian oversees our animal facility.

### Method details

### DNA constructs

All DNA constructs were cloned using the primers described in the “Oligonucleotides” section in the resource table. The primers described above for pCS2+/mNeon-VinculinR1049E were generated using NEBaseChanger (https://nebasechanger.neb.com/) to change CGG to GAA at sites 3142-3144 of Vinculin’s sequence (creating an arginine to glutamic acid mutation). Mutagenesis was achieved using a Q5 Site-Directed Mutagenesis Kit (New England BioLabs) following the manufacturer’s protocol. pCS2+/Vinculin was generated by PCR amplification of Vinculin from pCS2+/mNeon-Vinculin [47] and cloned into a digested pCS2+ vector (BamHI-HF and EcoRI-HF) via a Gibson reaction. Both constructs were verified by sequencing (GENEWIZ, South Plainfield, NJ and Plasmidsaurus, Eugene, OR).

### mRNA preparation and microinjections

To transcribe mRNA *in vitro*, all DNA constructs were linearized using NotI-HF, transcribed with the mMessage mMachine SP6 Transcription Kit (Invitrogen), and purified with the RNeasy Mini Kit (Qiagen). After purification, mRNA size was confirmed by running it on a 1% agarose gel with 1% bleach. mRNA was stored at -80°C until use.

In experiments without morpholino, a 5 nl volume was injected into the animal hemisphere of the embryos four times at either the 2-cell or 4-cell stage. Each 5 nl injection contained the following amount of mRNA or dye: pCS2+/Stargazin-GFPsilent-LOVPEP: 5 pg; 2xPDZ-YFP-LARG(DH): 2 pg; Halo-Vinculin: 10 pg; mNeon-Vinculin: 10 pg; LifeAct-RFP: 16 pg; SF9-mCherry: 74 pg; Angulin-1-3xGFP: 25 pg; Halo Janelia Fluor 646: 5 μM; Halo Janelia Fluor JFX650: 5 μM.

In experiments with morpholino, a 10 nl volume was injected into the animal hemisphere of the embryos four times at either the 2-cell or 4-cell stage. Each 10 nl injection contained the following amount of mRNA or dye, as well as morpholino or water (vehicle control): pCS2+/Stargazin-GFPsilent-LOVPEP: 5 pg; 2xPDZ-YFP-LARG(DH): 2 pg; mNeon-Vinculin: 10 pg; mNeon-VinculinR1049E: 10 pg; Vinculin: 10 pg; LifeAct-RFP: 16 pg; SF9-mCherry: 74 pg; Angulin-1-3xGFP: 25 pg; mNeon-Actin: 17 pg; Halo Janelia Fluor 646: 5 μM, Vinculin morpholino: 2.5 mM.

### Immunofluorescence staining

Paraformaldehyde (PFA) fixation was performed as follows for phosphomyosin, Vinculin, and F-actin immunofluorescence experiments: gastrula-stage embryos were placed in a mixture of 1.5% PFA, 0.25% glutaraldehyde, 0.2 % Triton X-100, and Alexa Fluor 568 phalloidin (1:100) in 0.88X MT buffer (800 mM K-PIPES, 5 mM EGTA, 1mM MgCl_2_, pH to 6.8) and allowed to fix on a shaker overnight at room temperature. Fixed embryos were quenched in 100 mM sodium borohydride in PBS on a shaker for 1 hour at room temperature. Embryos were then bisected to keep the animal cap and blocked with blocking solution (10% FBS, 5% DMSO, 0.1% NP-40 in 1X Tris-buffered Saline) overnight on a nutator at 4°C. The animal caps were then incubated overnight on a nutator at 4°C in the blocking solution with rabbit ɑ-phosphomyosin (1:100) and mouse ɑ-Vinculin (1:400). Next, they were washed three times in blocking solution and incubated overnight on a nutator at 4°C in the blocking solution with Goat Alexa Fluor 647 ɑ-rabbit (1:200), Goat Alexa Flour 488 ɑ-mouse (1:200), and Alexa Fluor 568 phalloidin (1:100). Embryos were washed and mounted in PBS before imaging.

Trichloroacetic acid (TCA) fixation was performed as follows for Angulin-1 immunofluorescence experiments: gastrula-stage embryos were placed in 2% TCA in PBS and allowed to fix for 2 hours on a shaker at room temperature. Fixed embryos were then bisected to keep the animal cap, blocked with blocking solution, permeabilized with 1% Triton X-100 in PBS for 20 minutes, permeabilized with 1X PBST (0.1% Triton X-100 in PBS) for 20 minutes, and blocked with blocking solution (5% FBS in 1X PBST) overnight at 4°C. The animal caps were then incubated overnight at 4°C in the blocking solution with rabbit ɑ-Angulin-1 (1:50). Next, they were washed three times in blocking solution and incubated overnight at 4°C in the blocking solution with Goat Alexa Fluor 647 ɑ-rabbit (1:200). Embryos were washed and mounted in PBS before imaging.

### Live imaging barrier assay

For ZnUMBA experiments, gastrula-stage embryos were injected one time in the blastocoel with 10 nl of a mixture of 100 µM CaCl_2_, 100 µM EDTA, and 1 mM FluoZin-3. 5 minutes post-injection, embryos were mounted in 1 mM ZnCl_2_ in 0.1X MMR before imaging [31, 32].

### Microscope image acquisition

All images were captured using an inverted Olympus FluoView 1000 confocal microscope with mFV-10-ASW software. Images were obtained with a supercorrected Plan Apo N 60XOSC objective (NA = 1.4, working distance = 0.12 mm). All live embryos were mounted in a chamber in a metal slide and held in place between two coverslips attached to the slide with vacuum grease.

#### Optogenetic stimulation and image acquisition

Time-lapse movies were acquired for Halo-Vinculin (with Janelia Fluor 646 or Janelia Fluor JX 650) and F-actin (LifeAct-RFP) by sequentially scanning the 8 apical Z-planes (step size of 0.37 μm) of a 512 x 512-pixel area (1.5X zoom) with a 559-nm laser and a 635-nm laser at 8 μs/pixel. During live imaging, simultaneous optogenetic stimulation was performed with the SIM scanner by creating a 512 x 512 region of interest and scanning the area with a 3% 405-nm laser at 2 µs/pixel. Videos were acquired by imaging without stimulation for 5 minutes (for a before stimulation baseline), followed by 5 minutes of simulation for 1 second every 20 seconds.

Time-lapse movies were acquired for F-actin (LifeAct-RFP) to measure morphological changes at TCJs in control and Vinculin KD embryos by scanning between 20-22 apical Z-planes (step size of 0.6 μm) of a 512 x 512-pixel area (1.5X zoom) with a 559-nm laser at 8 μs/pixel. During live imaging, simultaneous optogenetic stimulation was performed by creating a 512 x 512 region of interest and scanning the area with a 3% 405-nm laser at 2 µs/pixel. Videos were acquired by imaging without stimulation for 10 minutes (for a before stimulation baseline), followed by 10 minutes of simulation for 1 second every 20 seconds.

#### ATP treatment and image acquisition

Time-lapse movies were acquired for mNeon-Vinculin and F-actin (LifeAct-RFP) by sequentially scanning the 8 apical Z-planes (step size of 0.37 μm) of a 512 x 512-pixel area (3X zoom) with a 488-nm laser and a 559-nm laser at 10 μs/pixel. Embryos were mounted by sandwiching them between two coverslips, but only partially covering the hole with the top coverslip, creating an opening in the imaging chamber (**Fig 1B**). After 7-10 frames (for a pre-ATP baseline), 100 µl of 500 µM ATP in 0.1X MMR was added to the imaging chamber.

#### Fixed image acquisition

Fixed images were acquired by sequentially scanning the 15-20 apical Z-planes (step size of 0.37 μm) of a 512 x 512-pixel area (1.5X zoom) with the appropriate lasers (488-nm laser, 559- nm laser, and 635-nm laser) at 12.5 μs/pixel.

#### FRAP image acquisition

Time-lapse movies were acquired for FRAP experiments by scanning the Z-plane with the brightest signal of a 250 x 250-pixel area (2X zoom) with a 488-nm laser at 8 μs/pixel. Photobleaching was performed using the SIM scanner with the clip tornado function of the Olympus FV1000 mFV-10-ASW software. Videos were acquired by imaging for 3 frames (for a pre-bleach baseline) and then a 30% 405-nm laser was pulsed in a circular region that encompassed the junction and the neighboring cytosol (6 µm in diameter) for 600 msec (see **Fig S3A**).

#### ZnUMBA live imaging barrier assay acquisition

Time-lapse movies were acquired for the barrier assay by sequentially scanning the 8 apical Z-planes (step size of 0.37 μm) of a 320 x 320-pixel area (1.5X zoom) with a 488-nm laser (for FluoZin3 signal) and a 559-nm laser (LifeAct-RFP) at 8 μs/pixel. Imaging began immediately after submerging the embryo in 1 mM ZnCl_2_ in 0.1X MMR and continued for 30 minutes.

### Image processing and quantification

Except for the 3D projected image (**Fig 4B**), all images in the figures were max projected across each channel and were manually adjusted to show relevant features in Fiji. The 3D projected image was prepared using Imaris. Unless otherwise stated, all post-acquisition adjustments are consistent between control and Vinculin KD conditions within an experiment. If images were differentially adjusted between conditions, a scale bar with maximum and minimum values was added to each image. All quantification was performed using Fiji, normalization was performed using Excel, and statistical analysis was performed using GraphPad Prism.

#### Quantification of Vinculin intensity before and during optogenetic stimulation

Videos were sum projected, and for each embryo, one frame before RhoA activation and one frame during RhoA activation were chosen for quantification. The frames were selected to ensure Vinculin signal was in focus, and the during RhoA activation frame selected was the frame that displayed maximum contraction from RhoA activation. Using a 6 µm circular ROI, Halo-Vinculin signal was measured at seven TCJs before and during RhoA activation in each video. With the same ROI, three cytosolic measurements were taken and averaged. TCJ measurements were normalized to their corresponding average cytosolic signal. Finally, all TCJ measurements were normalized to the average of the TCJ measurements taken before RhoA activation to set the pre-stimulation baseline to 1. A paired t-test was performed to compare before and during RhoA activation Halo-Vinculin intensity.

#### Quantification of Vinculin intensity before and after ATP treatment

Videos were sum projected, and for each embryo, one frame pre-ATP addition and one frame post-ATP addition were chosen for quantification. The frames were selected to ensure Vinculin signal was in focus, and the post-ATP frame selected was the frame that displayed maximum contraction from ATP treatment. Using a 6 µm circular ROI, mNeon-Vinculin signal was measured at seven TCJs pre-and post-ATP treatment in each video. With the same ROI, three cytosolic measurements were taken and averaged. TCJ measurements were normalized to their corresponding average cytosolic signal. Finally, all TCJ measurements were normalized to the average of the TCJ measurements taken pre-ATP addition to set the pre-ATP treatment baseline to 1. A paired t-test was performed to compare pre-and post-ATP addition mNeon-Vinculin intensity.

#### Quantification of junctional phalloidin and phosphomyosin signal at TCJs

Images were max projected, and using a 6 µm circular ROI, phalloidin and phosphomyosin signal were measure at seven TCJs in each image. TCJ measurements were normalized to the highest signal measured and then normalized to set the baseline to 1. A one-way ANOVA test was performed to compare control, Vinculin KD, KD + WT, and KD + R1049E embryos.

#### Quantification of FRAP recovery curves

All FRAP videos were max projected. For TCJ FRAP experiments, the intensity of the bleached region of interest and a nearby reference TCJ was quantified in all frames using a circular ROI (2.90 μm diameter for mNeon-Actin; 1.45 μm diameter for Angulin-3xGFP). For BCJ FRAP experiments, the intensity of the bleached region of interest and a nearby reference BCJ was quantified in all frames using a line ROI (3.5 μm long, width 2). Each measurement was taken in triplicate and then averaged. The intensities were normalized by the following formula: I_norm_(t) = (I_ref pre_ / I_ref_(t)) * (I_frap_(t) / I_frap pre_). Where I_ref_ is the intensity of the reference junction, I_frap_ is the intensity of the bleached junction, (t) is the specific time point, and *_pre_* is the average intensity pre bleach. The normalized intensities were then constrained between 1 and 0 with the following formula: I_norm1_(t) = (I_norm_(t) - I_norm_(t_bleach_)) / (I_norm pre_ - I_norm_(t_bleach_)). Where I_norm_ is the normalized value from the first formula, (t) is the specific time point, (t_bleach_)is the bleached, and *_pre_* is the average intensity pre bleach time point. The I_norm1_ values were plotted over time on GraphPad Prism and fit to a double exponential curve in order to extrapolate the mobile fraction and fast/slow t_1/2_. An unpaired t-test with Welch’s correction was done to compare control and Vinculin KD conditions.

#### Quantification of distance TCJs dip before and during RhoA activation

For each embryo, one frame before RhoA activation and one frame during RhoA activation were chosen for quantification. The frames were selected to ensure the LifeAct signal was in focus, and the during RhoA activation frame selected was the frame that exhibited maximum contraction from RhoA activation. Using the XZ views in Fiji of 7 different TCJs per embryo, the distance TCJs dip was determined by measuring the distance between the apical surface and the start of the TCJ signal (**Fig 4C**). If the TCJ signal was *above* the apical surface, the distance was recorded as zero. A paired t-test was performed to compare before and during RhoA activation.

#### Quantification of barrier function

All ZnUMBA videos were max projected, and the file names were coded to blind the quantification. Only videos that were at least 25 minutes long were quantified; for videos over 25 minutes, only the first 25 minutes were quantified. Criteria for leaky TCJs were as follows: a TCJ was only counted one time throughout the video regardless of whether it leaked repeatedly or not, leaks had to be at least 5 µm in diameter to be counted, the leaks had to be at TCJs (we did not count multicellular junctions that had 4 or more junctions intersecting), and we did not count leaks at TCJs that were at the cleavage furrow of dividing cells. All TCJs in a given video were counted. The percentage of leaky TCJs was determined by dividing the number of leaky TCJs by the number of total TCJs. An unpaired t-test was performed to compare control and Vinculin KD conditions.

## Acknowledgments

We thank all the members of the Miller lab for helpful discussions and feedback on this research. This work was funded by the National Institutes of Health (R01GM112794 to A.L.M.). L.vdG. is supported by an American Heart Association Predoctoral Fellowship.

## Author contributions

Conceptualization, L.vdG. and A.L.M.; Methodology, L.vdG and A.L.M.; Formal Analysis, L.vdG. and J.I.; Investigation, L.vdG., J.I., and K.K.; Resources, L.vdG. and A.L.M.; Writing – Original Draft, L.vdG.; Writing – Review & Editing, L.vdG, J.I., K.K., and A.L.M.; Visualization, L.vdG; Supervision, A.L.M.; Funding Acquisition, L.vdG. and A.L.M.

## Video Legends

**Video S1: Vinculin is mechanosensitively recruited to cell-cell junctions in response to optogenetic RhoA activation, related to Figure 1**.

Halo-Vinculin (with JF646) is recruited to cell-cell junctions upon 405-nm light stimulation to activate RhoA-mediated contraction (area and duration of light stimulation indicated by magenta box). Time shown in minutes:seconds. Playback at 4 fps.

**Video S2: Vinculin is mechanosensitively recruited to cell-cell junctions in response to extracellular addition of ATP, related to Figure 1**.

mNeon-Vinculin is recruited to cell-cell junctions upon extracellular addition of ATP (ATP addition indicated by magenta box). Time shown in minutes:seconds. Playback at 5 fps.

**Video S3: Vinculin maintains Angulin-1 protein stability at TCJs, related to Figure 3**.

Videos showing Angulin-1-3xGFP at TCJs in control (top) and Vinculin KD (bottom) embryos before, during and after photobleaching. Images are shown using the FIRE LUT, adjusted in the same way for each image. Bleaching is indicated by yellow circles. Time shown in minutes:seconds. Playback at 20 fps.

**Video S4: Vinculin maintains TCJ morphology in response to optogenetic RhoA activation, related to Figure 4**.

The first video shows control embryos expressing LifeAct-RFP. The left panel shows full Z-projection, and the right panel shows apical Z-projection. Cells contract in response to optogenetic RhoA activation (indicated by magenta box), but TCJs remain intact. The second video shows Vinculin KD embryos expressing LifeAct-RFP. The left panel shows full Z-projection, and the right panel shows apical Z-projection. In response to optogenetic RhoA activation (indicated by magenta box), TCJs are disrupted. Time shown in minutes:seconds. Playback at 5 fps.

**Video S5: Vinculin maintains barrier function at TCJs, related to Figure 5**.

Video showing FluoZin3 signal in control (left) and Vinculin KD (right) embryos. Images are shown using the FIRE LUT, adjusted in the same way for each image. Arrows indicate leaks at TCJs. Time shown in minutes:seconds. Playback at 6 fps.

