## Supplemental Material for "Mechanosensitive recruitment of Vinculin maintains junction integrity and barrier function at epithelial tricellular junctions"

#### Supplemental Figure Legends

**Figure S1: Actomyosin organization in response to increased tension, related to Figure 1.**

**A)** Live confocal images before and during RhoA activation of embryos expressing an F-actin probe (LifeAct-RFP) and (**A'**) a myosin probe (SF9-mCherry) and Vinculin (Halo-Vinculin with JF646).

**B)** Live confocal images before and after addition of extracellular ATP of embryos expressing an F-actin probe (LifeAct-RFP), (**B'**) a myosin probe (SF9-mCherry) and Vinculin (mNeon-Vinculin), and (**B"**) the tTJ protein Angulin-1 (Angulin-1-3xGFP).

**Figure S2: Vinculin is mechanosensitively recruited to BCJs, related to Figure 1.**

**A)** Quantification of Halo-Vinculin intensity at BCJs before and during RhoA activation mediated by optogenetic stimulation. Statistics, paired t-test; n = 3 experiments, 9 embryos, 54 BCJs; p ≤ 0.0001 (****). Violin plots show the median (dashed line) and the 25$^{th}$ and 75$^{th}$ quartiles (dotted lines).

**B)** Quantification of mNeon-Vinculin intensity at BCJs before and after extracellular ATP addition. Statistics, paired t-test; n = 3 experiments, 5 embryos, 29 BCJs; p ≤ 0.0001 (****). Violin plots show the median (dashed line) and the 25$^{th}$ and 75$^{th}$ quartiles (dotted lines).

**Figure S3: Validation of Vinculin knockdown, related to Figures 2 and 3.**

**A)** Schematic of the custom-designed Gene Tools Vinculin morpholino (Vinculin MO) which binds to the 5' UTR of endogenous Vinculin mRNA, thus blocking translation.

**B)** *Left:* Percent identity and similarity of full-length Vinculin comparing Human (*Homo sapiens*), Mouse (*Mus musculus*), and Frog (*Xenopus laevis*) sequences. *Right:* Sequence alignment of the Vinculin tail between Human, Mouse, and Frog, highlighting that R1049 is conserved between the three species.

**C)** Fixed confocal images of control, Vinculin KD, KD + WT, and KD + R1049E embryos that were stained for Vinculin (α-Vinculin) and F- actin (phalloidin).

**D)** Quantification of junctional intensity of α-Vinculin. Statistics, one way ANOVA; n = 2 experiments, control = 17 embryos, Vinculin KD = 12 embryos, KD + WT = 1123 embryos, KD + R1049E = 12 embryos; p-values > 0.05 (ns), ≤ 0.05 (*), ≤ 0.01 (**), ≤ 0.0001 (****). Violin plots show the median (dashed line) and the 25$^{th}$ and 75$^{th}$ quartiles (dotted lines).

**E)** Quantification of cell size. Statistics, one way ANOVA; n = 2 experiments control = 17 embryos, Vinculin KD = 12 embryos, KD + WT = 1123 embryos, KD + R1049E = 12 embryos; p-values ≤

0.05 (*), ≤ 0.0001 (****). Violin plots show the median (dashed line) and the 25th and 75th quartiles (dotted lines).

**F)** Western blot showing endogenous protein levels of Angulin-1, actin, and tubulin in control, Vinculin KD, and KD + WT embryos.

**Figure S4: FRAP analysis and measured mobile fraction and recovery rate values, related to Figures 2 and 3.**

**A)** Schematic showing bleached TCJ and BCJ areas and measured TCJ and BCJ areas for FRAP experiments.

**B)** Table of measured mobile fractions and recovery rate values for mNeon-Actin FRAP at BCJs and TCJs. Values were calculated from recovery curves shown in **Fig 2E**. Statistics comparing control and Vinculin KD, unpaired t-test; p-value ≤ 0.0001 (****); SDs are indicated.

**C)** Table of measured mobile fractions and recovery rate values for Angulin-1-3xGFP FRAP at TCJs. Values were calculated from recovery curves shown in **Fig 3D**. Statistics comparing control and Vinculin KD, unpaired t-test; p-values ≤ 0.01 (**), ≤ 0.0001 (****); SDs are indicated.

**Supplemental Methods**

*Quantification of junctional Vinculin signal*

For Vinculin KD validation via staining, junctional Vinculin signal was measured by creating a mask of junctional signal in FIJI. For all images, maximum projection was used. The phalloidin channel was used to create the mask as follows: the image was duplicated to create two copies (Images A and B). A Gaussian blur with a radius of 15 was applied to Image B to remove noise. Using the image calculator, Image B was subtracted from Image A to create Image C. A Gaussian blur with a radius of 3 was applied to Image C to detect continuous junctions. Then, thresholding was applied to Image C implementing the Huang method. The signal was then dilated one time. Any cytosolic signal in Image C was manually deleted using the poly selection tool and the image was inverted. Image C was then subtracted from the maximum projection of the Vinculin channel until all cytosolic signal was removed from the Vinculin channel. The mean intensity of the Vinculin channel was then measured and recorded, and a one-way ANOVA test was performed to compare control, Vinculin KD, KD + WT, KD + R1049E embryos.

*Western blotting*

Gastrula-stage embryos were washed in chilled PHEME lysis buffer (60 mM K-PIPES, 25 mM HEPES, 10 mM, EGTA, 2 mM $MgCl_2$, pH 7.0 using KOH). The embryos were then homogenized using a pellet pestle (7495211500, DWK Life Sciences) in PHEME lysis buffer ensuring each group had the same number of embryos. The lysed embryos were transferred to chilled spin columns (344718, Beckman Coulter) and centrifuged at 20,817 x g for 5 minutes at 4ºC. The cytoplasmic layer was then removed using a 1 mL syringe and a 27 G ½ inch needle to puncture the side of the spin columns and placed in a new chilled tube. 1/3 volume of 6X SDS was then added to the lysate, and the samples were boiled for 10 minutes. The prepared lysates were separated by SDS-PAGE (4-20%, Mini Protean TGX Precast Protein gel (4561094, Bio-Rad)), transferred to nitrocellulose using Trans-Blot Turbo Transfer System (Bio-Rad) for 7 mins at 25 V, and blocked using StartingBlock (PBS) Blocking Buffer (37538, Thermo Scientific). Primary antibodies were diluted in blocking buffer in incubated overnight at 4ºC. The membrane was washed three times with TBST before being incubated in secondary antibodies diluted in blocking buffer for two hours at room temperature. The membrane was then washed three times in TBST, one time in PBS, and developed using Pierce ELC Western Blotting Substrate (32209, Thermo Scientific). After imaging, the membrane was stripped and re-probed using Restore Western Blot Stripping Buffer (PI21059, Thermo Scientific). Primary antibodies were used at the following concentrations: rabbit anti-Angulin-1 (custom), 1:500; mouse anti-actin (A4700, Sigma),

1:500; mouse anti-tubulin (T9026, Sigma), 1:10,000. Secondary antibodies were used at the following concentrations: anti-mouse HRP (PRW4021, Promega) and anti-rabbit HRP (PRW4011, Promega).

*Cell size quantification*

Cell size measurements were performed by creating a mask for phalloidin immunofluorescent images. For all images, maximum projection was used. The phalloidin channel was used to create the mask as follows: the image was duplicated to create two copies (Images A and B). A Gaussian blur with a radius of 15 was applied to Image B to remove noise. Using the image calculator, Image B was subtracted from Image A to create Image C. A Gaussian blur with a radius of 3 was applied to Image C to detect continuous junctions. Then, thresholding was applied to Image C, adjusting the min/max to maximize junction continuity. The junctions were then dilated until the junctions were continuous. The image was then skeletonized and dilated 1 time. The image was inverted, and cell size was measured using “Analyze particles” with the following parameters: size: 20 – infinity, show outlines, display results, exclude on edges, include holes. The outlines were used to confirm the mask resembles the cell outlines, and the areas were recorded. Finally, a one-way ANOVA test was performed to compare control, Vinculin KD, KD + WT, KD + R1049E embryos.

#### Figure S1: Actomyosin organization in response to increased tension, related to Figure 1

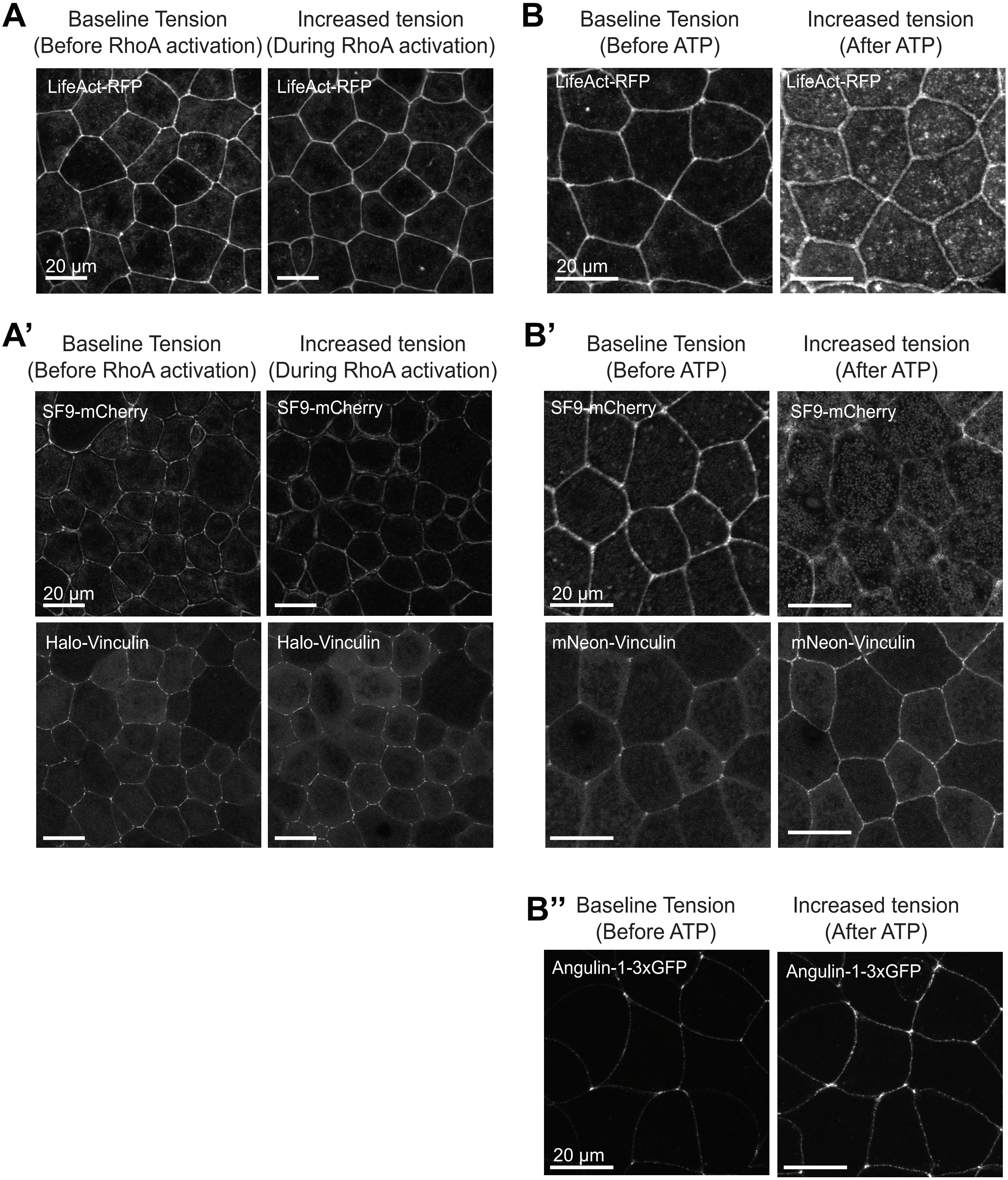

#### Figure S2: Vinculin is mechanosensitively recruited to BCJs, related to Figure 1

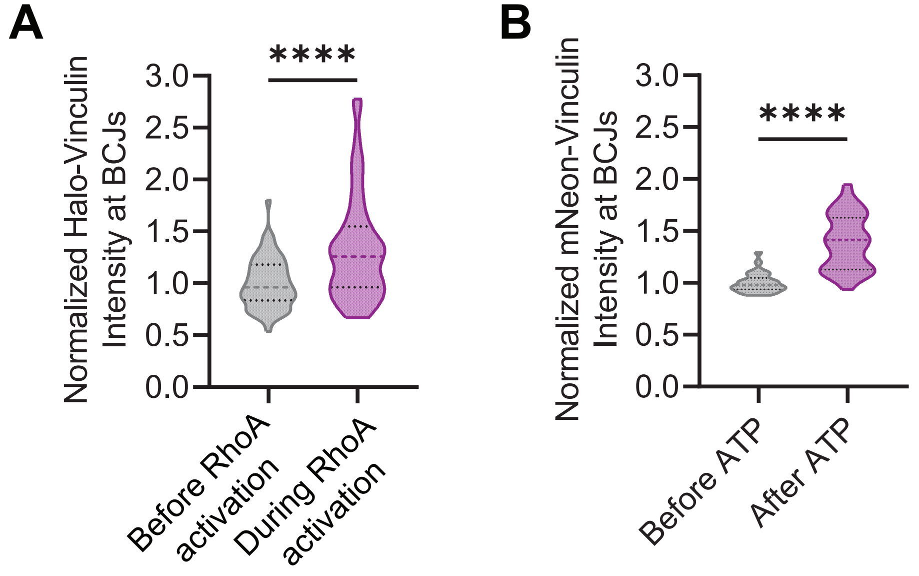

### Figure S3: Validation of Vinculin knockdown, related to Figures 2 and 3

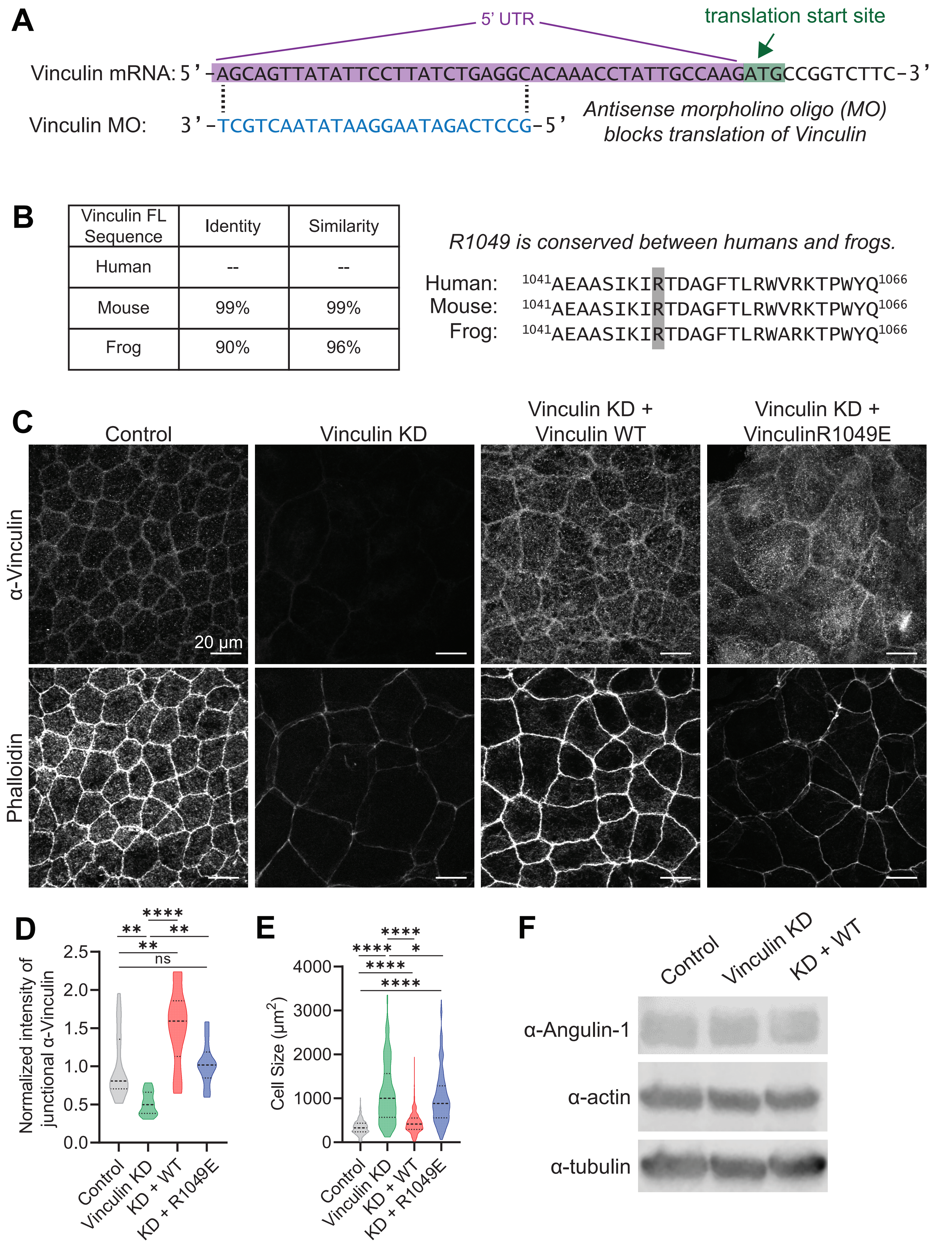

### Figure S4: FRAP analysis and measured mobile fraction and recovery rate values, related to Figures 2 and 3

A

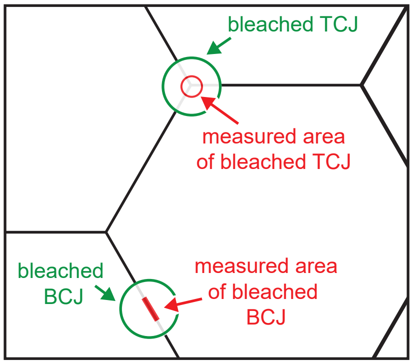


B

| | Actin | mobile fraction | fast $t_{1/2}$ | slow $t_{1/2}$ |
|---|---|---|---|---|
| BCJ | control | 86.1 ± 0.81% | 0.73 ± 0.17 s | 17.1 ± 0.86 s |
| BCJ | Vinculin KD | 83.8 ± 0.81%**** | 1.24 ± 0.28 s | 14.1 ± 1.02 s |
| TCJ | control | 86.5 ± 0.63% | 0.73 ± 0.12 s | 16.5 ± 0.76 s |
| TCJ | Vinculin KD | 95.7 ± 0.45%**** | 0.67 ± 0.17 s**** | 11.6 ± 0.40 s |

C

| | Angulin-1 | mobile fraction | fast $t_{1/2}$ | slow $t_{1/2}$ |
|---|---|---|---|---|
| TCJ | control | 49.4 ± 0.04% | 4.72 ± 2.90 s | 83.9 ± 14.58 s |
| TCJ | Vinculin KD | 63.1 ± 0.01%**** | 1.43 ± 2.17 s**** | 38.1 ± 2.80 s** |
